# Heterologous expression of *PtAAS1* reveals the metabolic potential of the common plant metabolite phenylacetaldehyde for auxin synthesis *in planta*

**DOI:** 10.1101/2022.12.15.520544

**Authors:** Jan Günther, Rayko Halitschke, Jonathan Gershenzon, Meike Burow

## Abstract

Aromatic aldehydes and amines are common plant metabolites involved in several specialized metabolite biosynthesis pathways. Recently, we showed that the aromatic aldehyde synthase PtAAS1 and the aromatic amino acid decarboxylase PtAADC1 contribute to the herbivory-induced formation of volatile 2-phenylethanol and its glucoside 2-phenylethyl-β-D-glucopyranoside in *Populus trichocarpa*. To gain insights into alternative metabolic fates of phenylacetaldehyde and 2-phenylethylamine beyond alcohol and alcohol glucoside formation, we expressed *PtAAS1* and *PtAADC1* heterologously in *Nicotiana benthamiana* and analyzed plant extracts using untargeted LC-qTOF-MS analysis. While the metabolomes of *PtAADC1*-expressing plants did not significantly differ from those of control plants, expression of *PtAAS1* resulted in the accumulation of phenylacetic acid (PAA) and PAA-amino acid conjugates, identified as PAA-aspartate and PAA-glutamate. Moreover, targeted LC-MS/MS analysis showed that *PtAAS1*-expressing plants accumulated significant amounts of free PAA. The measurement of PAA and PAA-Asp in undamaged and herbivory-damaged poplar leaves revealed significantly induced accumulation of PAA-Asp while levels of free PAA remained unaltered by herbivore treatment. Sequence comparisons and transcriptome analysis showed that members of a small gene family comprising five putative auxin-amido synthetase *GH3* genes potentially involved in the conjugation of auxins like PAA with amino acids were significantly upregulated upon herbivory in *P. trichocarpa* leaves. Overall, our data indicates that phenylacetaldehyde generated by poplar PtAAS1 serves as a hub metabolite linking the biosynthesis of volatile, non-volatile herbivory-induced specialized metabolites, and phytohormones, suggesting that growth and defense are balanced on a metabolic level.

## Introduction

Plant specialized metabolites mediate plant responses to different biotic conditions and are generated by a plethora of biosynthetic pathways. These pathways can be initiated by key enzymes like cytochrome P450 enzymes (Irmisch et al., 2013; Sørensen et al., 2018), aminotransferases (Wang and Maeda, 2018), and group II pyridoxal phosphate (PLP)-dependent enzymes (Facchini et al., 2000). The latter comprise decarboxylase and aldehyde synthase enzymes that are involved in the biosynthesis of aromatic amino acid-derived specialized metabolites like benzylisoquinoline alkaloids (Facchini et al., 2000), monoterpene indole alkaloids (O’Connor and Maresh, 2006), hydroxy cinnamic acid amides (Facchini et al., 2002), and phenylpropanoids in plants (Torrens-spence et al., 2018; Günther et al., 2019). For example, poplar trees under herbivore attack biosynthesize the volatile 2-phenylethanol and its glucoside 2-phenylethyl-β-D-glucopyranoside via separate biosynthetic pathways (Günther et al., 2019). The formation of these metabolites can be initiated by the closely related group II PLP-dependent enzymes, PtAAS1 and PtAADC1. The direct reaction products of AAS and AADC enzymatic reactions phenylacetaldehyde and 2-phenylethylamine have been shown to contribute to the biosynthesis of different plant metabolites, respectively (Sekimoto et al., 1998; Facchini et al., 2000; Facchini et al., 2002; Torrens-spence et al., 2018). Previously, it has been shown that 2-phenylethylamine could be transformed to phenylacetaldehyde in subsequent biosynthetic steps in planta (Boatright et al., 2004; Tieman et al., 2006). The aromatic aldehyde phenylacetaldehyde has further been proposed to contribute to the biosynthesis of the auxin phenylacetic acid (Cook and Ross, 2016; Cook et al., 2016).

Auxins are plant hormones of paramount impact on plant development and growth. Although indole-3-acetic acid (IAA) is the most studied amongst this group of phytohormones, other natural auxins like phenylacetic acid (PAA), indole-3-butyric acid, indole-3-propionic acid and 4-chloroindole-3-acetic acid have been discovered (Abe et al., 1974; Fries and Iwasaki, 1976; Ludwig-Müller, 2011; Simon and Petrášek, 2011). Recent progress has led to an increased interest in the biosynthesis and physiology of the auxin PAA (Enders and Strader, 2016; Zhao, 2018). In comparison, IAA and PAA show distinctly different characteristics concerning their transport and distribution within the plant (Sugawara et al., 2015; Aoi et al., 2020). Both auxins target similar responsive elements, whereas a lower concentration of IAA in comparison to the concentration of PAA is sufficient to trigger these responses (Wightman and Lighty, 1982; Simon and Petrášek, 2011). Cellular concentrations of auxins are variable, and their developmental output is highly dependent on the local biosynthesis and local concentrations of these auxins (Wang et al., 2015; Zheng et al., 2016). Free auxin concentrations can be rapidly altered by auxin-amido synthetase Gretchen Hagen 3 (GH3) enzymes (Westfall et al., 2016). These enzymes belong to a large family of auxin amino acid synthetases that accept IAA as well as PAA and conjugate these substrates with different amino acids (Staswick et al., 2005; Sugawara et al., 2015). Conjugation has been shown to interfere with both signaling and transport of free auxins and thereby modulate signaling in plant development (Ljung et al., 2002; Ludwig-Müller, 2011; Zheng et al., 2016). Furthermore, specific fluctuations in auxin biosynthesis and transport trigger different developmental changes in the context of adaptation to stress (Grieneisen et al., 2007; Brumos et al., 2018; Zhao, 2018; Blakeslee et al., 2019). Generally, auxins like IAA and PAA can be biosynthesized via separate biosynthetic pathways within the plant kingdom (Pollmann et al., 2006; Mano and Nemoto, 2012; Zhao, 2014; Cook and Ross, 2016). To date, much is known about the biosynthetic pathways leading to the formation of IAA in *Arabidopsis thaliana* (Mashiguchi et al., 2011; Zhao, 2014; Enders and Strader, 2016). Initial steps of this biosynthetic network comprise the formation of indole-3-acetaldehyde and tryptamine (Supplemental Figure 1; reviewed in (Mano and Nemoto, 2012; Zhao, 2014)). In further biosynthetic steps, these intermediates can be converted to the corresponding auxin IAA. Similarly, the biosynthesis of PAA can be initiated by separate pathways leading to the formation phenylacetaldehyde (Kaminaga et al., 2006; Gutensohn et al., 2011; Günther et al., 2019) and 2-phenylethylamine (Tieman et al., 2006; Günther et al., 2019). In subsequent biosynthetic steps, these pathways might ultimately lead to the formation of PAA (Supplemental Figure 2; (Sekimoto et al., 1998)).

In this study, we show that the key enzyme for generation of the herbivory-induced metabolites phenylacetaldehyde, 2-phenylethanol and 2-phenylethyl-β-D-glucopyranoside in poplar lead to stimulated biosynthesis of the auxin PAA as well as PAA conjugates *in planta*.

## Results and Discussion

### Expression of *PtAAS1* and *PtAADC1* alters the accumulation of phenolic metabolites in *N. benthamiana*

We expressed *PtAAS1* and *PtAADC1* in leaves of *N. benthamiana* and quantified the aromatic amino acid substrates and aromatic amine products of AADC1 as well as the indirect reaction product of AAS1, 2-phenylethyl-β-D-glucopyranoside, via LC-MS/MS. As previously shown, levels of aromatic amines (Supplemental Figure 3) and 2-phenylethyl-β-D-glucopyranoside (Supplemental Figure 4) were increased upon *PtAADC1* and *PtAAS1* expression, respectively (Günther et al., 2019).

To investigate other potential metabolic alterations in *PtAAS1*- and *PtAADC1*-expressing *N. benthamiana* leaves, we performed untargeted LC-qTOF-MS analysis. The expression of *PtAAS1* and *PtAADC1* resulted in different metabolite profiles in comparison to wild type and *eGFP*-expressing control plants (Figure 1; Supplemental Table 1). The expression of *PtAAS1* and *PtAADC1* resulted in the differential accumulation of metabolites with a higher number of significantly up- or downregulated metabolites in the *PtAAS1*-expressing plants (Figure 1). Two candidate metabolites were exclusively present in *PtAAS1*-expressing lines and were identified as conjugates of phenylacetic acid with aspartate (PAA-Asp) and glutamate (PAA-Glu) via LC-qTOF-MS as described recently (Westfall et al., 2016; Aoi et al., 2020). Additionally, we observed characteristic insource fragmentation patterns of PAA-Asp and PAA-Glu in negative ionization mode, respectively (Supplemental Figure 5). Several other phenolic compounds were detected but could not be identified based on the fragmentation pattern (Supplemental Table 1).

**Figure 1:**
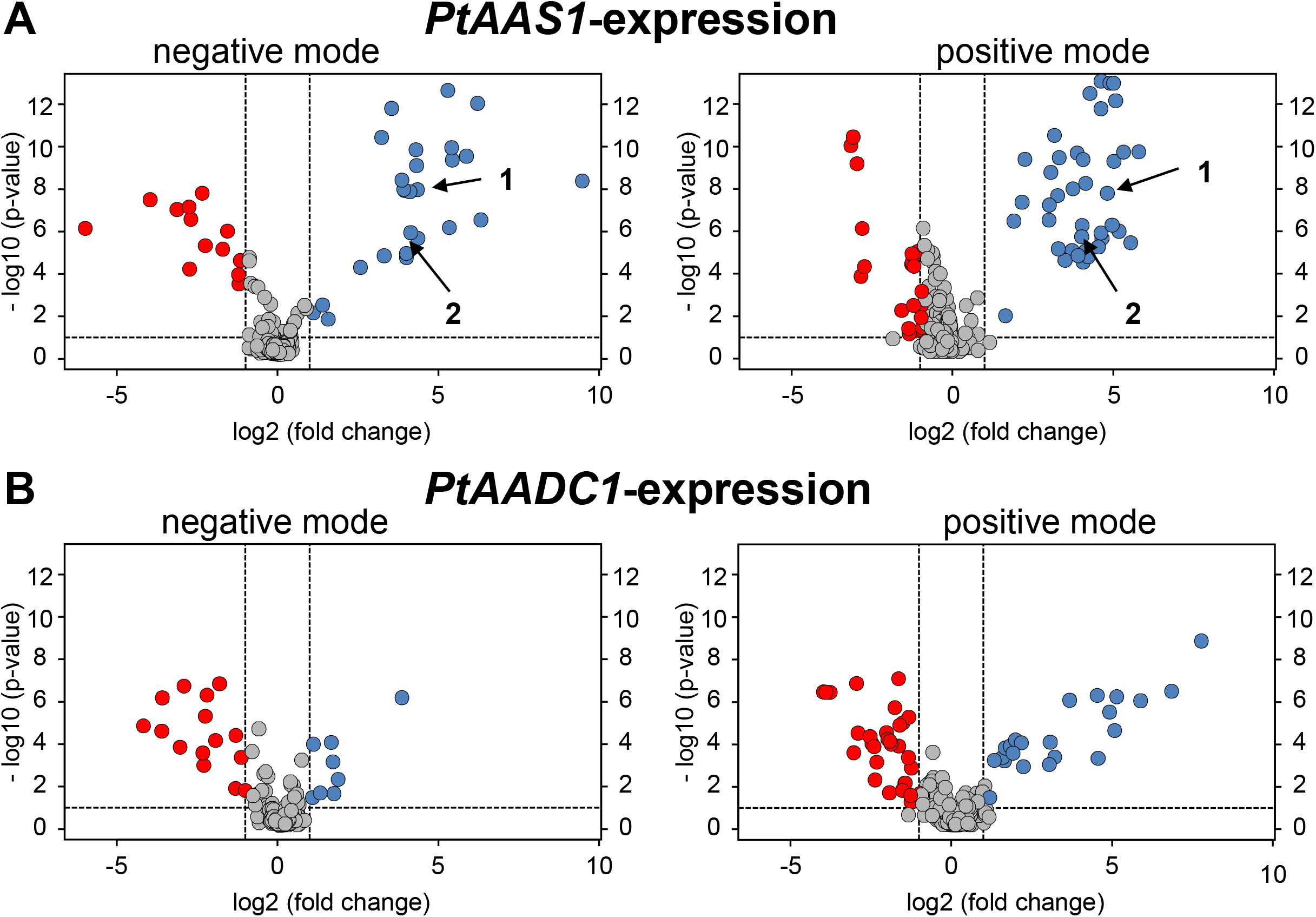
Untargeted LC-qTOF-MS reveals significantly altered metabolites in *PtAAS1*-and *PtAADC1-*expressing *N. benthamiana* leaves. Volcano plots of normalized LC-qToF-MS analysis of significantly upregulated (blue) and downregulated (red) metabolites in (A) *PtAAS1*- and (B) *PTAADC1*-expressing *N. benthamiana plants* in comparison to *eGFP*-expressing control plants (n = 6).

Expression of *PtAAS1* and the concomitant accumulation of auxin-conjugates pointed towards a conversion of the PtAAS1 reaction product phenylacetaldehyde to PAA-Asp and PAA-Glu in further metabolic steps in a heterologous plant system. It has been shown recently that aromatic aldehyde synthases generate aldehydes from corresponding aromatic amino acids and thereby initiate to the formation of aromatic alcohols and alcohol glucosides (Torrens-Spence et al., 2018; Günther et al., 2019). Additionally, it has been suggested that aromatic aldehydes might also contribute to the formation of plant auxins (Sekimoto et al., 1998).

In addition to the accumulation of 2-phenylethyl-β-D-glucopyranoside, *PtAAS1*-expressing *N. benthamiana* plants also accumulated the auxin PAA in leaves (Figure 2). In comparison to reports in *Arabidopsis thaliana* rosette leaves, which were shown to contain up to 500 pmol/g fresh weight (∼70ng/g fresh weight) of endogenous PAA (Sugawara et al., 2015), the amounts of up to 700 ng/g fresh weight in leaves of *PtAAS1*-expressing *N. benthamiana* plants (Figure 2) are unphysiologically high. These high concentration of the indirect PtAAS1 product PAA might have allowed highly promiscuous endogenous enzymes to accept this metabolite as a substrate (Moghe and Last, 2015). Such enzymes might be employed for detoxification of toxic intermediates within specialized metabolism (Sirikantaramas et al., 2008). Indeed, plant auxins can be inactivated or detoxified via esterification with amino acids (Woodward and Bartel, 2005; Korasick et al., 2013). Nevertheless, our results in *N. benthamiana* illustrate that PtAAS1 activity can lead to the formation of the phenylalanine-derived auxin PAA in a heterologous plant system.

**Figure 2:**
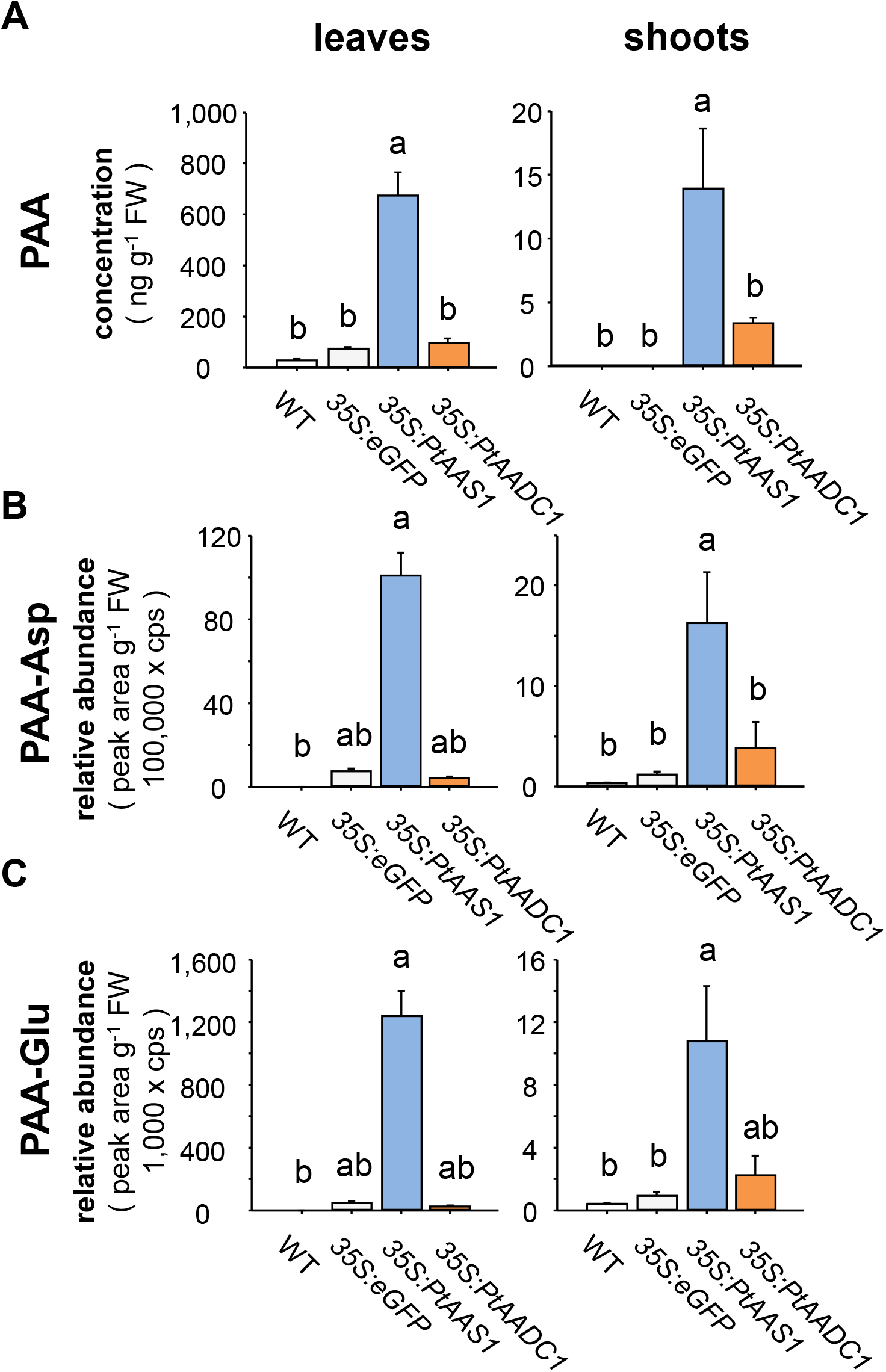
Expression of *PtAAS1* results in increased levels of the auxin PAA, and its conjugates PAA-Asp and PAA-Glu in *N. benthamiana* leaves and shoots. *N. benthamiana* leaves expressing poplar *PtAAS1* accumulate high amounts of (A) PAA, (B) PAA-Glu and (C) PAA-Asp in leaves and shoots. Different letters above each bar indicates statistically significant differences (Kruskal-Wallis One Way ANOVA) and are based on the following Tukey (shoots) or Dunńs test (leaves): PAA_leaves_ (H = 19.607, P ≤ 0.001); PAA_shoot_ (H = 12.275, P = 0.006); PAA-Asp_leaves_ (H = 20.747, P ≤ 0.001); PAA-Asp_shoot_ (H = 19.127, P ≤ 0.001); PAA-Glu_leaves_ (H = 19.924, P ≤ 0.001); PAA-Glu_shoot_ (H = 15.647, P = 0.001). Means + SE are shown (n = 6). FW, fresh weight.

### Expression of phenylacetaldehyde-generating PtAAS1 leads to the accumulation of PAA- and IAA-conjugates in *N. benthamiana*

We further investigated the accumulation of PtAAS1-derived auxin metabolites in different tissues of *N. benthamiana* plants expressing *PtAAS1*. In order to further characterize the function of PtAAS1 and PtAADC1 we developed targeted analyses to quantify the reaction products and the auxin derivatives.

The auxin conjugates PAA-Asp and PAA-Glu accumulated in leaves and shoots (Figure 2). Furthermore, we detected these auxin conjugates in roots, however, their levels in roots were not significantly increased in *PtAAS1*-expressing plants in comparison to wild type and *eGFP*-expression control plants (Supplemental Figure 6). It has been reported that *A. thaliana* plants with an increased PAA biosynthesis also showed an increased accumulation of PAA-Asp, PAA-Glu and IAA-aspartate conjugate (IAA-Asp; (Aoi et al., 2020)). In line with these findings, we measured significantly increased levels of IAA-Asp in leaves and shoots of *PtAAS1*-expressing plants (Supplemental Figure 7), highlighting that the increased auxin biosynthesis is accompanied by increased conversion of both auxins IAA and PAA into their respective conjugates (Mashiguchi et al., 2011; Sugawara et al., 2015; Aoi et al., 2020). Auxin conjugation with amino acids occurs in plant tissue upon the increase of auxin biosynthesis or accumulation (Ludwig-Müller, 2011) and regulates the concentration of free, active auxin, to mitigate the developmental effects of increased auxin biosynthesis (Zhao et al., 2001; Mashiguchi et al., 2011; Bunney et al., 2017).

We identified that *PtAAS1* contributes to the formation of the auxin PAA and its conjugates PAA-Asp and PAA-Glu in *N. benthamiana* leaves (Figure 1; 2). It has been shown recently that expression of *CYP79A2* resulted in similar increase of PAA, PAA-Asp and PAA-Glu in Arabidopsis (Aoi et al., 2020). Our results indicate that the expression of *PtAAS1* in *N. benthamiana* results in the increased biosynthesis of PAA and confirms the results of Aoi and colleagues that levels of other active auxins might be reduced within the plant by means of conjugation of the active forms to the inactive conjugates (Aoi et al., 2020). Furthermore, we could show that the PAA conjugates accumulate in the leaf tissue as well as in adjacent shoot and root tissue (Figure 2; Supplemental Figure 6). These results allow for speculation of a directional transport of PAA conjugates that were biosynthesized in the leaves as reviewed recently (Leyser, 2018).

Plant auxins are involved in various stages of plant development and defense (Grieneisen et al., 2007; Brumos et al., 2018; Günther et al., 2018; Zhao, 2018; Blakeslee et al., 2019). Local auxin concentration is strictly regulated within plants by means of transport, degradation and conjugation (Ljung et al., 2002; Staswick et al., 2005; Ludwig-Müller, 2011; Korasick et al., 2013; Sugawara et al., 2015; Zheng et al., 2016) and external application of auxins results in the increased accumulation of auxin conjugates. In this study, we induced high accumulation of the auxin PAA in a heterologous plant system via the expression of the phenylacetaldehyde-generating *PtAAS1* (Figure 1; 2). The resulting accumulation of the corresponding PAA conjugates suggests that the pool size of free auxins is strictly regulated, at least partially by conversion into inactive conjugates and possibly transport of these conjugates.

Taken together, *PtAAS1* contributes to the formation of PAA and PAA conjugates *in planta*. Additionally, as we also detected increased levels of IAA-Asp, our results provide additional evidence for the recently described crosstalk of PAA with IAA through coordinated conjugation of both free auxins (Aoi et al., 2020).

### The auxin conjugate PAA-Asp and putative *GH3* transcripts accumulate in herbivory-induced poplar leaves

We next tested whether herbivory-induced *P. trichocarpa* leaves with increased *PtAAS1* transcript levels show accumulation of PAA and PAA-Asp. We incubated *P. trichocarpa* trees with the herbivore *Lymantria dispar* caterpillars and quantified PAA and PAA-Asp in the leaves. Notably, the accumulation of PAA was unaltered in comparison to control leaves, whereas PAA-Asp was significantly increased upon herbivory (Figure 3). It has been previously shown that increased auxin biosynthesis can be accompanied by the stimulated expression of auxin responsive elements like *GH3* and *Aux/IAA* genes as well as the reduction of *SAUR* gene expression (Hagen and Guilfoyle, 2002). To evaluate whether transcripts of these gene families are upregulated in *P. trichocarpa* leaves challenged by herbivorous enemies, we screened for putative *Aux/IAA*, *GH3* and *SAUR* genes in our in-house transcriptome dataset. Amongst the family of 14 *GH3* genes identified in the poplar transcriptome, five candidates were significantly induced in response to herbivory (Figure 3). Phylogenetic relationships of these herbivory-induced transcript suggests that these genes might indeed encode GH3 enzymes that catalyze the conjugation of the auxin PAA with amino acids (Supplemental Figure 8; (Staswick et al., 2005; Böttcher et al., 2011; Peat et al., 2012; Yu et al., 2018)). The *Aux/IAA* gene family in poplar consists of 15 putative members, two of which were significantly upregulated in herbivory-induced poplar leaves (Supplemental Figure 9). No transcripts of the 104-membered putative *SAUR* gene family were differentially expressed (Supplemental Table 2). These results suggest that upon herbivory, putative *GH3* transcripts accumulate in poplar. The corresponding GH3 enzymes might lead to the formation of PAA-Asp in herbivory-induced poplar leaves. At the time of harvest, 24 hours after the start of the herbivore treatment, the PAA concentration had most likely returned to basal levels, whereas the level of the conjugation product was still increased.

**Figure 3:**
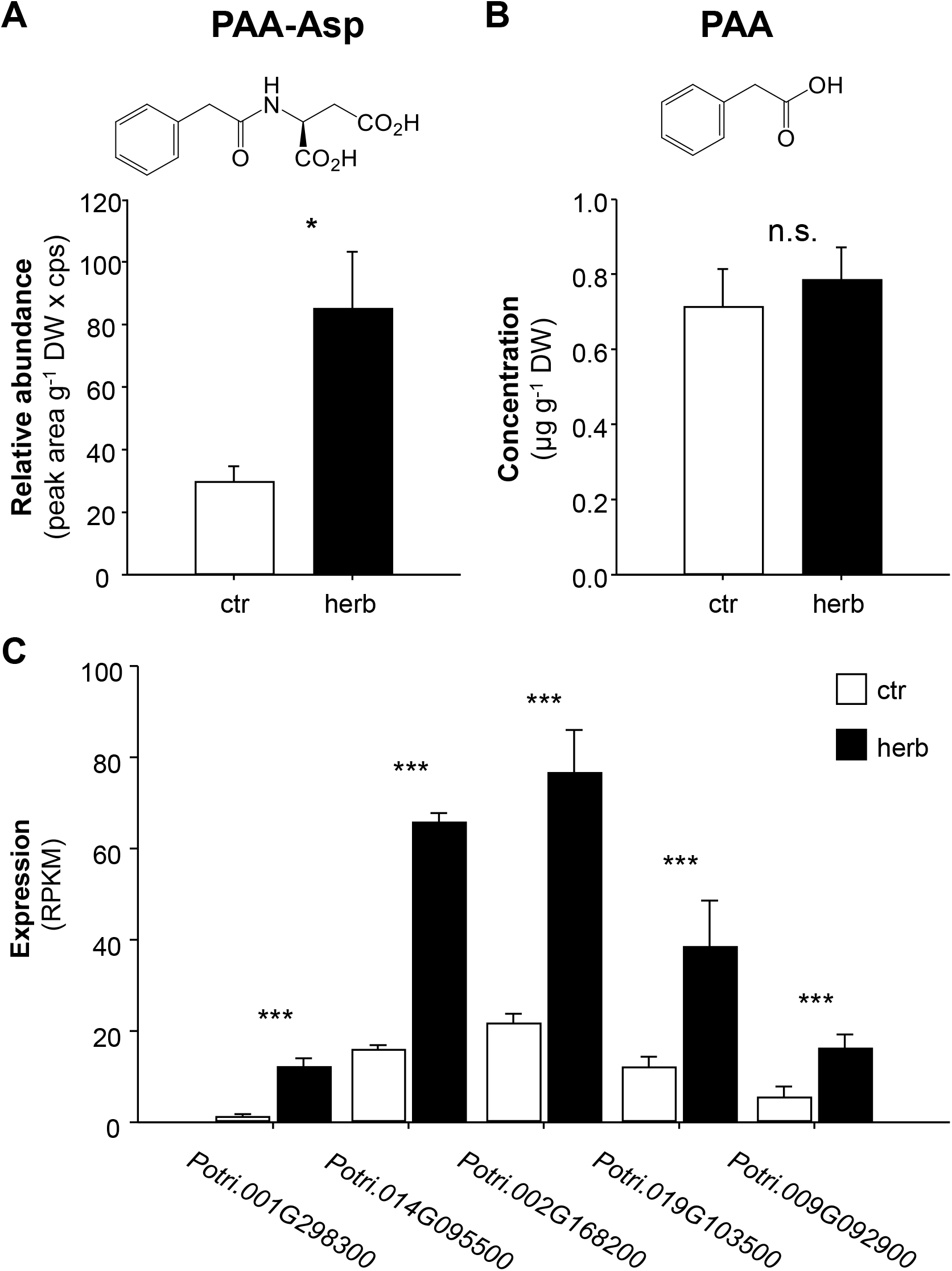
Auxin, auxin conjugate and putative auxin-amido synthetase GH3 transcripts accumulate in herbivore-damaged leaves of *Populus trichocarpa*. Accumulations of PAA-Asp (A) and phenylacetic acid (B), were analyzed in *L. dispar* damaged (herb) and undamaged control (ctr) leaves of *Populus trichocarpa* via LC-MS/MS. Asterisks indicate statistical significance in Student’s t-test or in Mann-Whitney Rank Sum Tests. PAA (P = 0.608, t = -0.522); PAA-Asp (P = 0.011, t = -2.816). Putative auxin-amido synthetase *GH3* Gene expression (C) in herbivore-damaged and undamaged leaves was analyzed by Illumina HiSeq sequencing. Expression was normalized to RPKM. Significant differences in EDGE tests are visualized by asterisks. Means + SE are shown (n = 4). *Potri.001G298300* (P = 2.22705E-10, weighted difference (WD) = 1.71922E-05); *Potri.014G095500* (P = 4.04229E-20, WD = 7.9334E-05); *Potri.002G168200* (P = 1.01033E-12, WD = 8.83786E-05); *Potri.019G103500* (P = 1.39832E-05, WD = 4.27809E-05); *Potri.009G092900* (P = 5.97467E-05, WD = 1.72578E-05). Means + SE are shown (n = 10). DW, dry weight. n.s. – not significant.

Taken together, our results highlight a potential unprecedented role of PtAAS1 in the biosynthesis of the auxin PAA *in planta*. We hypothesize that the expression of *PtAAS1* in *N. benthamiana* leaves leads to an increased metabolic flow to generate high amounts of volatile 2-phenylethanol, 2-phenylethyl-β-D-glucopyranoside (Günther et al., 2019), PAA and PAA-conjugates in response to an increased biosynthesis of the hub metabolite phenylacetaldehyde. As suggested recently, plant specialized metabolites might be integrated into regulation of plant signaling, growth and development (Erb and Kliebenstein, 2020).

Several studies revealed that the expression of key biosynthetic enzymes that initiate the biosynthesis of aromatic amino acid-derived specialized metabolites resulted in altered auxin phenotypes and chemotypes (Bak and Feyereisen, 2001; Bak et al., 2001; Irmisch et al., 2015; Günther et al., 2018; Perez et al., 2021; Perez et al., 2022). In Arabidopsis, another link between specialized metabolism and auxin signaling might be established by indole glucosinolate hydrolysis products with high affinity to the major auxin receptor Transport Inhibitor Response 1 (TIR1), which could result in competitive binding and thereby feed into the auxin signaling cascade (Vik et al., 2018). Therefore, it may not only be auxin itself but also structural analogues like indole glucosinolates and their hydrolysis products that trigger auxin-induced developmental effects. Additionally, it has been recently reported, that other non-aromatic glucosinolate catabolites are able to stimulate auxin-like phenotypes in Arabidopsis roots (Katz et al., 2015; Katz et al., 2020). Similarly, the maize defense compounds benzoxazolinones might contribute to auxin-induced growth through the interference with auxin perception (Hoshi-Sakoda et al., 1994), as maize CYP79A enzymes contribute to the formation of phenylalanine-derived defense compounds as well as to the formation of the corresponding auxin PAA in a heterologous plant system (Irmisch et al., 2015). Finally, the most recent study of Aoi and colleagues showed that the expression of *CYP79A2* that leads to the formation of (*E*/*Z*)-phenylacetaldoxime in Arabidopsis resulted in effects similar to those of increased auxin biosynthesis and auxin conjugation (Aoi et al., 2020). Several different pathways classified as specialized metabolism appear to have evolved to not only mediate plant biotic interaction, but also provide regulatory input to auxin signaling. Increased expression of biosynthetic enzymes involved in the biosynthesis of amino acid-derived specialized metabolites might directly influence the homeostasis of the corresponding auxins.

In summary, the poplar aromatic aldehyde synthase PtAAS1 contributes to the herbivory-induced formation of volatile 2-phenylethanol and 2-phenylethyl-β-D-glucopyranoside and additionally to the formation of the auxin PAA and auxin-derived conjugates in a heterologous plant system. We show that the conjugate of PAA-Asp accumulates upon herbivory in poplar leaves, suggesting that herbivory-induced expression of *PtAAS1* might contribute to PAA biosynthesis and might stimulate PAA signaling and metabolism in poplar. We conclude that the biosynthesis of the hub metabolite phenylacetaldehyde is of paramount importance for the generation of the auxin PAA and represents an additional pathway for the formation of the auxin PAA, expanding the metabolic network of the convergent biosynthesis of this auxin *in planta* (Figure 4). We unraveled unprecedented aspects of the biosynthesis of the auxin PAA in a heterologous plant system as well as in response to herbivory in *P. trichocarpa* leaves. Therefore, the phenylacetaldehyde hub metabolite represents a metabolic link between volatile, non-volatile herbivory-induced specialized metabolites, and phytohormones, suggesting that both growth and defense are balanced on a metabolic level. Further research needs to be aimed at elucidating and understanding the plant physiological responses following the increased PAA biosynthesis and conjugation upon herbivory.

**Figure 4:**
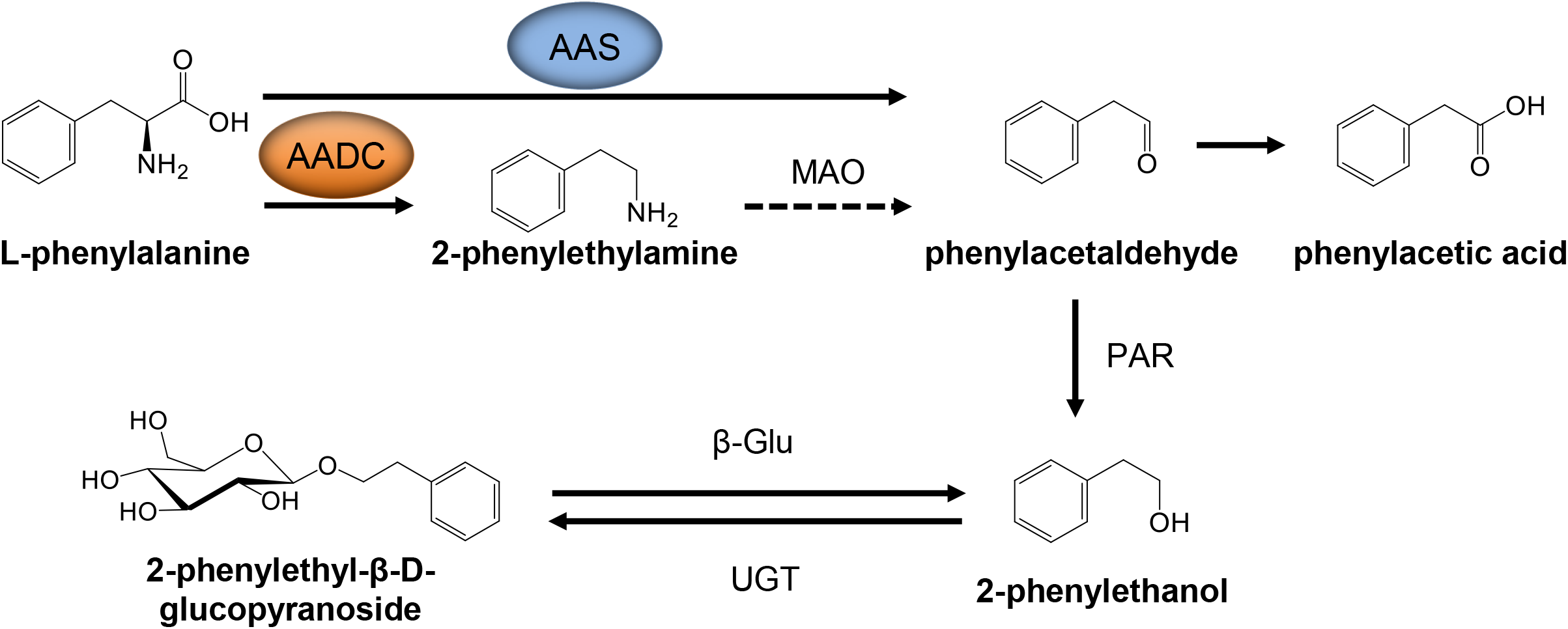
Proposed pathways for the convergent biosynthesis of PAA in planta. Convergent biosynthesis of 2-phenylethanol that can be initiated by AAS and AADC enzymes has been described. The initiation of the formation of phenylacetaldehyde as common substrate of 2-phenylethanol might also serve as substrate for the biosynthesis of the auxin PAA. Respective enzymes have been elucidated *in planta*. AADC, aromatic amino acid decarboxylase; AAS, aromatic aldehyde synthase; MAO, monoamine oxidase; PAR, phenylacetaldehyde reductase; UGT, UDP-glucosyl transferase; β-Glu, β-glucosidase. Dashed arrow, enzymes uncharacterized *in planta*. Solid lines, enzymes characterized *in planta*.

## Materials and Methods

### Plant material and treatment

*Populus trichocarpa* (genotype Muhle Larsen) trees were grown, *Lymantria dispar* herbivory was induced and all leaves (including midrib) and shoots (stem and petiole) were harvested. Roots were cleared of soil by washing in a fresh water bath, dried with a paper towel. All plant samples were frozen in liquid nitrogen immediately after harvesting. Plant samples were stored at -80 °C until further processing as described earlier (Günther et al., 2019). Agrobacterium-mediated expression of target genes in *N. benthamiana* was performed as described (Günther et al., 2019). Three days after transformation, plants were placed under mild direct light (LED 40%) for three more days.

### LC-qTOF-MS analysis of *N. benthamiana* methanol extracts

Methanol extracts (10:1 v/w) of *N. benthamiana* leaves were analyzed on an Ultimate 3000 UHPLC equipped with an Acclaim column (150 mm × 2.1 mm, particle size 2.2 µm) and connected to an IMPACT II UHR-Q-TOF-MS system (Bruker Daltonics) following a previously described program in positive and negative ionization mode (He et al., 2019). Raw data files were analyzed using Bruker Compass DataAnalysis software version 4.3. Metabolomic differences of extracts were analyzed via MetaboScape 4.0 (Supplemental Table 1). Furthermore, untargeted MS data was normalized and visualized in volcano plots via XCMS (Tautenhahn et al., 2012; Gowda et al., 2014; Rinehart et al., 2014; Benton et al., 2015; Johnson et al., 2016).

### LC-MS/MS analysis of plant methanol extracts

Metabolites were extracted from ground plant material (*P. trichocarpa* or *N. benthamiana*) with methanol (10:1 v/w). Analytes were separated using an Agilent 1200 HPLC system on a Zorbax Eclipse XDB-C18 column (5034.6 mm, 1.8 µm; Agilent Technologies). HPLC parameters are given in Supplemental Table 3. The HPLC was coupled to an API-6500 tandem mass spectrometer (Sciex) equipped with a turbospray ion source (ion spray voltage, 4500 eV; turbo gas temperature, 700 °C; nebulizing gas, 60 p.s.i.; curtain gas, 40 p.s.i.; heating gas, 60 p.s.i.; collision gas, 2 p.s.i.). Multiple reaction monitoring (MRM) was used to monitor a parent ion → product ion reactions given in Supplemental Table 4. The identification of PAA and IAA conjugates was performed according to LC-MS/MS fragmentation patterns as described in (Irmisch et al., 2013; Sugawara et al., 2015; Westfall et al., 2016; Günther et al., 2018; Günther et al., 2019). Relative quantification was based on the relative abundance in measured extracts based on counts per second in standardized measurement conditions. Identification and quantification of PAA, 2-phenylethylamine, tyramine, tryptamine, 2-phenylethyl-β-D-glucopyranoside, tyrosine, tryptophan and phenylalanine was performed with authentic, commercially available standards (Supplemental Table 5).

### Statistics

Statistical analysis was carried out as described in the figure legends. Student’s t tests, Mann-Whitney Rank Sum tests, Kruskal-Wallis one-way analysis of variance (ANOVA), Dunńs tests and Tukey tests were performed with the software SigmaPlot 14.0 (Systat Software). EDGE tests for analysis of RNA-Seq datasets were performed with CLC Genomics Workbench (Qiagen Informatics) as described earlier (Günther et al., 2019).

### RNA extraction, cDNA synthesis and RNA-Seq analysis

Poplar leaf RNA extraction, cDNA synthesis and RNA-Seq analysis were carried out as described (Günther et al., 2019). For the identification of putative *Aux/IAA*, *GH3* and *SAUR* genes in the *P. trichocarpa* genome (Tuskan et al., 2006), the transcriptome annotations as mapped to the poplar gene model version 3.0 provided by Phytozome (https://phytozome.jgi.doe.gov/pz/portal.html) were used for identification of members of the *Aux/IAA*, *GH3* and *SAUR* gene families. Candidates of the *Aux/IAA* and *GH3* gene family identified as herbivory-induced with above average fold change and p-values below P = 0.05 were selected for visualization (Figure 3; Supplemental Figure 9; Supplemental Table 2). Total RPKM counts of all control and all herbivore-induced treatments were summed for the estimate of total transcript differences within the *SAUR* gene expression (Supplemental Table 2).

### Phylogenetic analysis

Evolutionary analyses were conducted in MEGA X (Kumar et al., 2018)(Kumar et al., 2018). Coding sequences were retrieved from Phytozome (https://phytozome.jgi.doe.gov/pz/portal.html) and a multiple codon sequence alignment was performed via the guidance 2 server (Landan and Graur, 2008; Penn et al., 2010; Sela et al., 2015). The evolutionary history was inferred by using the Maximum Likelihood method based on the General Time Reversible model (Nei and Kumar, 2000). Initial trees were obtained automatically by applying Neighbor-Join and BioNJ algorithms to a matrix of pairwise distances estimated using the Maximum Composite Likelihood (MCL) approach, and then selecting the topology with superior log likelihood value. A discrete Gamma distribution was used to model evolutionary rate differences among sites (5 categories (+G, parameter = 1.5416)). The rate variation model allowed for sites to be evolutionarily invariable ([+I], 13.41% sites).

## Supporting information

Supplemental infromation

## Author Contributions

J.Gü. and J.Ge. designed research. J.Gü., carried out the experimental work, analyzed data and wrote the manuscript. R.H. analyzed samples via untargeted LC-qToF-MS. J.Ge. and M.B. contributed to and finalized the manuscript. All authors read and approved the final manuscript.

## Acknowledgments

We appreciate the helpful comments and suggestions of Tobias G. Köllner on the manuscript. We thank Tamara Krügel, Danny Kessler, and all the MPI-CE gardeners for their help with rearing the poplar and *Nicotiana benthamiana* plants. D. Werck-Reichhart, Strasbourg, France, is thanked for kindly providing the pCAMBIA vectors and for the nice introduction to the USER cloning system.

---

**Supplemental Figure 1:**
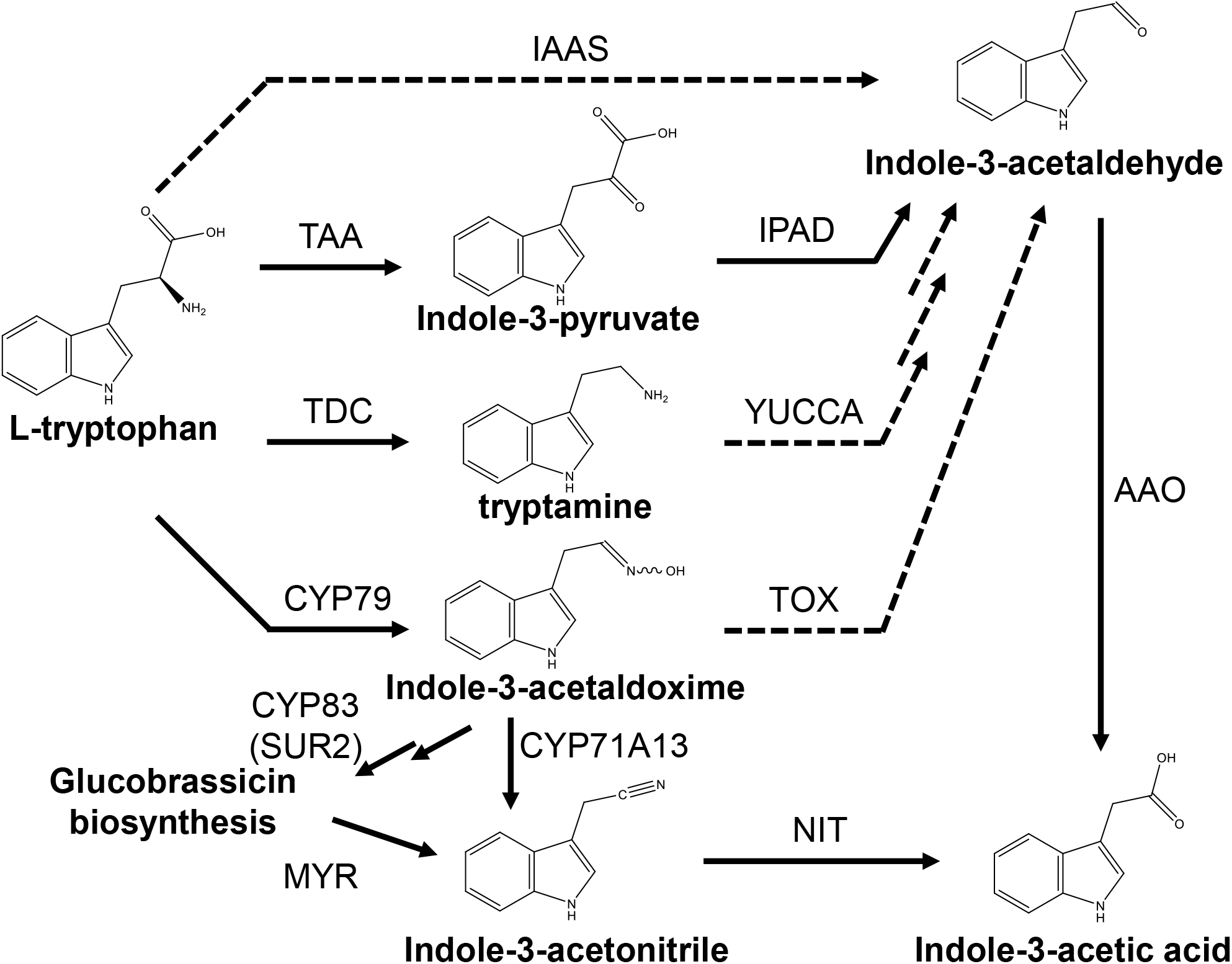
Proposed, simplified pathways for the biosynthesis of indole-3-acetic acid in plants. TAA, tryptophan aminotransferase; TDC, tryptophan decarboxylase; CYP79, cytochrome P450 family 79 enzyme; IAAS, indole-3-acetaldehyde synthase; IPA-DC, Indole-3-pyruvic acid decarboxylase; TOX, transoximase; AAO, aromatic aldehyde synthase; YUCCA, flavin monooxygenase-like enzyme; MYR, myrosinase; CYP83, cytochrome P450 family 83 enzyme; IPAD, indole-3-pyruvic acid decarboxylase. Dashed arrows, enzymes not characterized in plants; solid arrows, enzymes characterized in plants.

**Supplemental Figure 2:**
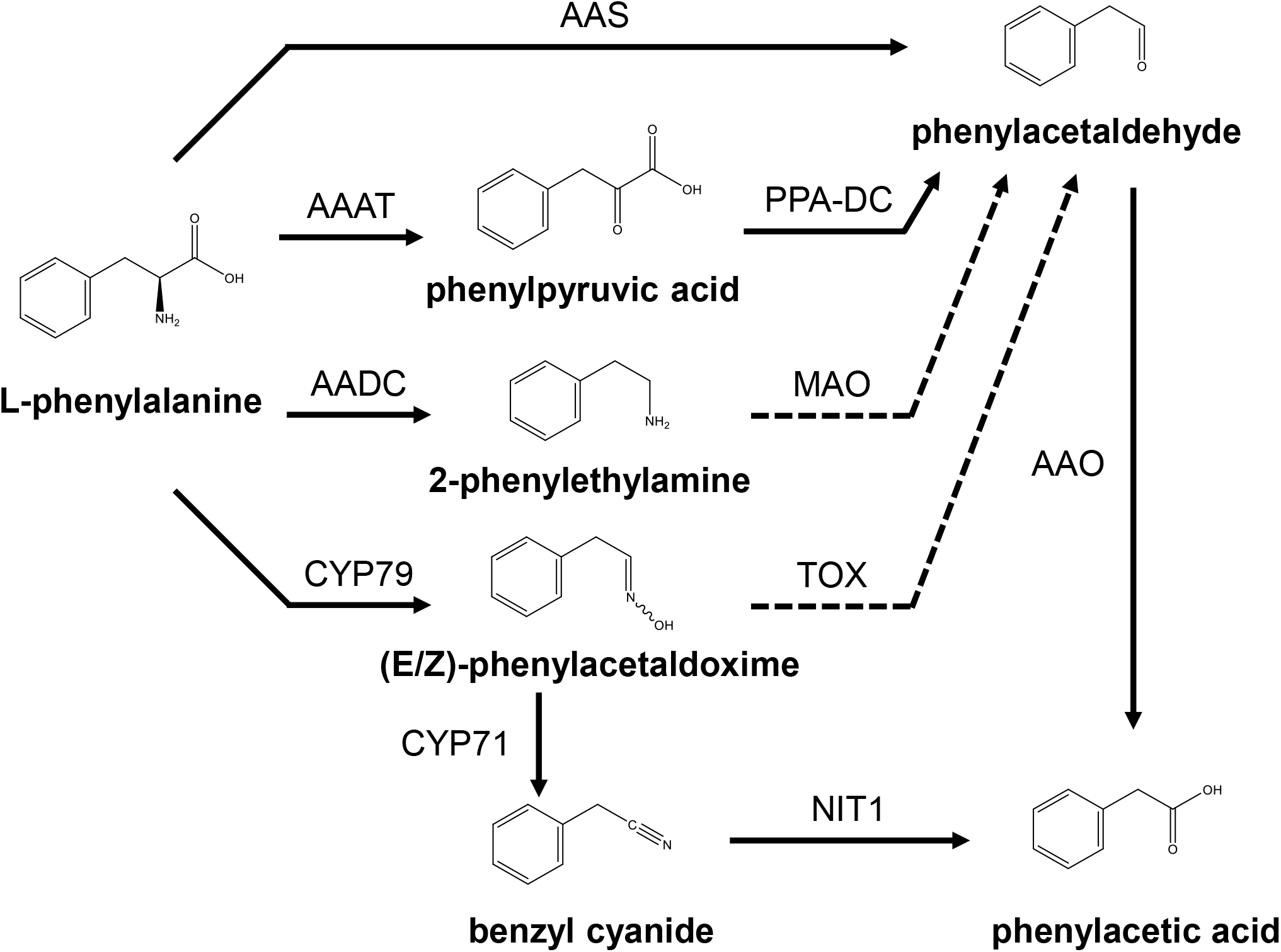
Proposed pathways for the biosynthesis of phenylacetic acid in plants. AAAT, aromatic amino acid transaminase; AADC, aromatic amino acid decarboxylase; CYP79, cytochrome P450 family 79 enzyme; PAAS, phenylacetaldehyde synthase; PPA-DC, phenylpyruvic acid decarboxylase; MAO, monoamine oxidase; TOX, transoximase; PAR, phenylacetaldehyde reductase; UGT, UDP-glucosyl transferase; β-Glu, β-glucosidase. Dashed line, enzymes not characterized in plants; solid line, enzymes characterized in plants.

**Supplemental Figure 3:**
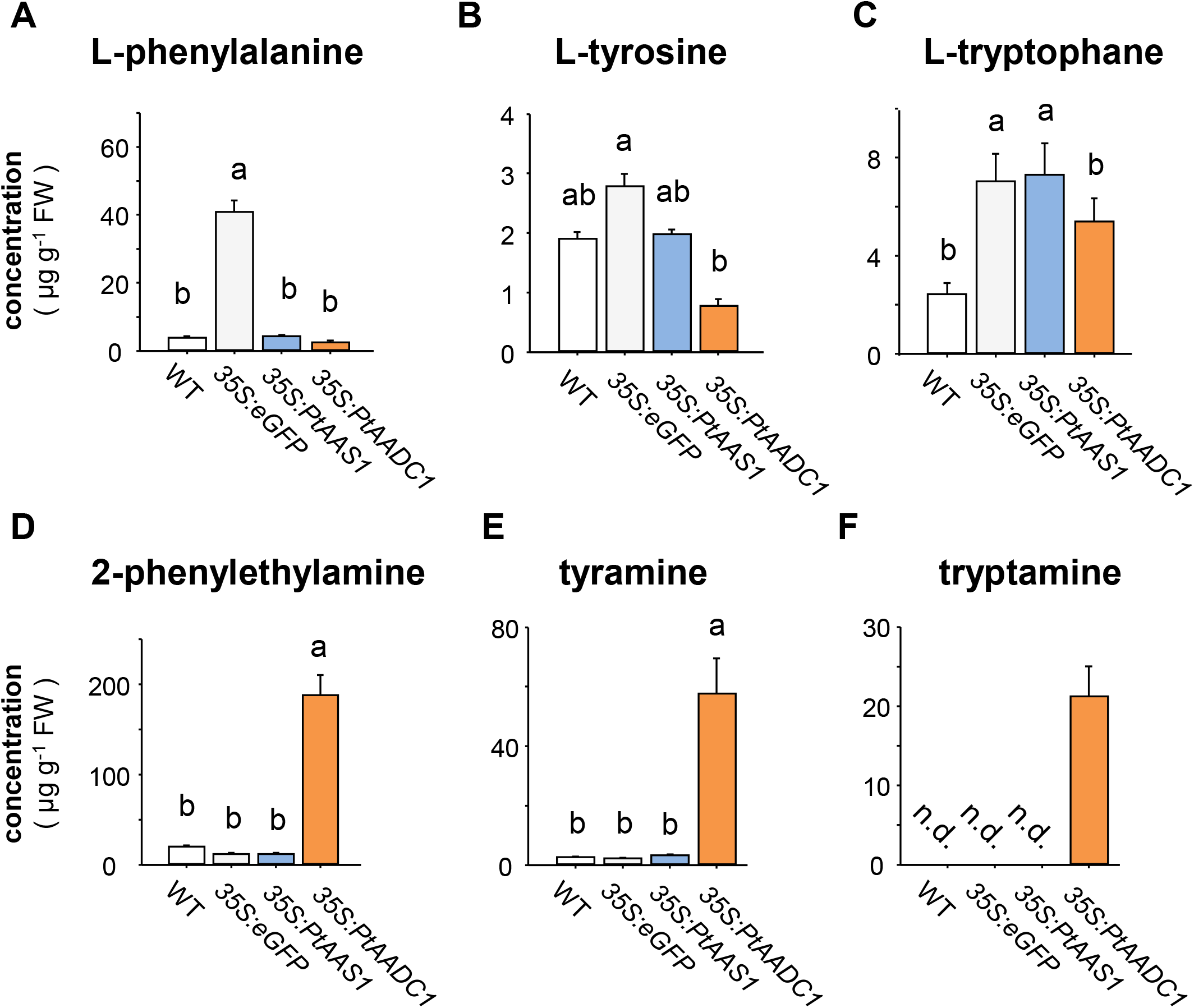
Expression of *PtAADC1* results in decreased aromatic amino acid substrate pools (A-C) and accumulation of aromatic amine products (D-F) in *N. benthamiana* leaves. The expression of poplar *PtAADC1* leads to the depletion of the aromatic amino acid substrates L-phenylalanine (A), L-tyrosine (B), and L-tryptophane (C) in expressing leaves. Accordingly, the corresponding enzymatic reaction products phenylethylamine (D), tyramine (E), and tryptamine (F) accumulate in *N. benthamiana* leaves, respectively. Different letters above each bar indicate statistically significant differences in Kruskal-Wallis One Way ANOVA and are based on the following Tukey test. Phe (H = 16.58, P ≤ 0.001); Tyr (F = 32.883, P ≤ 0.001); Trp (F = 73.043, P ≤ 0.001); PEA (H = 19.167, P ≤ 0.001); TyrA (H = 17.34, P ≤ 0.001). Means + SE are shown (n = 6). FW, fresh weight. n.d., not detected.

**Supplemental Figure 4:**
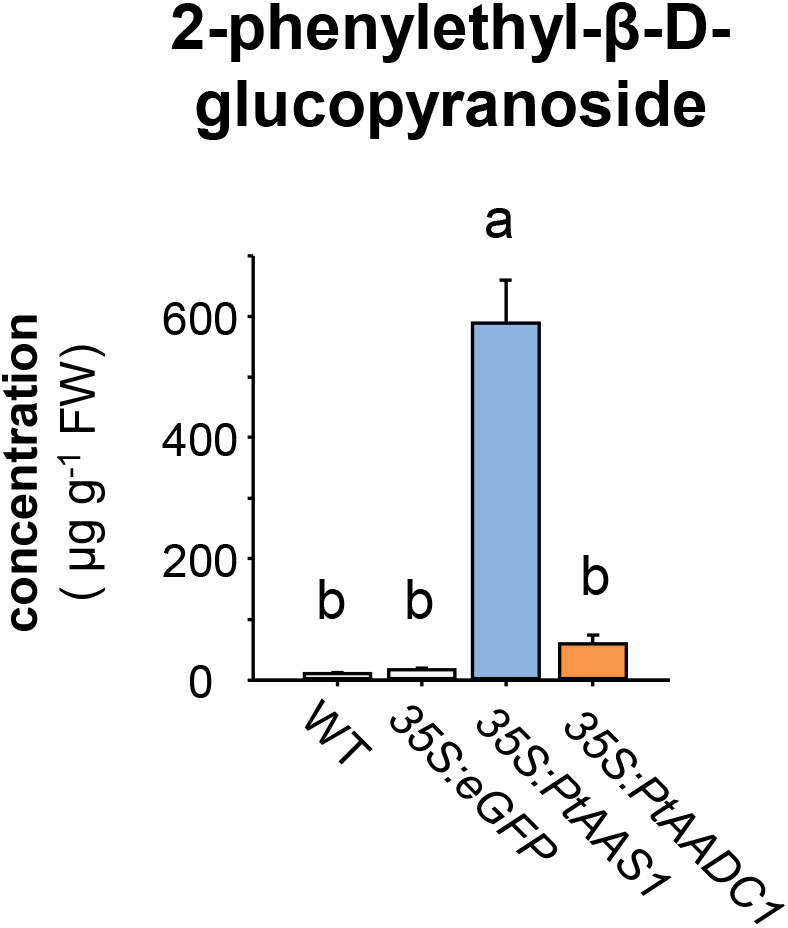
Expression of *PtAADC1* and *PtAAS1* results in the increased accumulation of 2-phenylethyl-β-D-glucopyranoside in *N. benthamiana* leaves. *N. benthamiana* plants expressing *eGFP*, *PtAAS1, PtAADC1* and wild type plants were grown for 5 days post inoculation as described (Günther et al., 2019). The accumulation of 2-PEG in *N. benthamiana* leaves was analyzed via LC-MS/MS. Different letters above each bar indicate statistically significant differences in Kruskal-Wallis One Way ANOVA and Tukey test. 2-PEG (H = 20.24, P ≤ 0.001). Means + SE are shown (n = 6). FW, fresh weight.

**Supplemental Figure 5:**
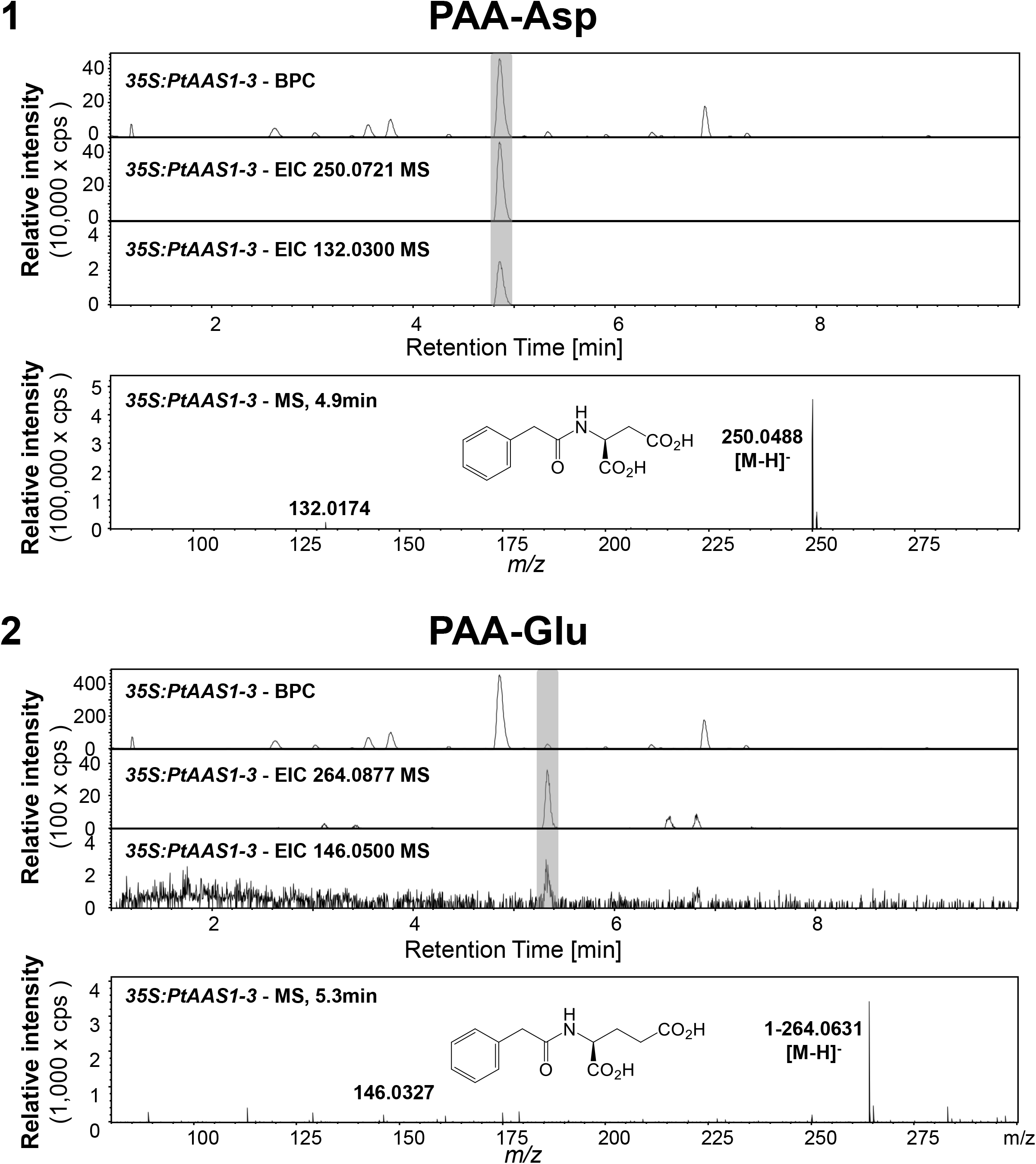
LC-qToF-MS analysis of PAA conjugates in negative ionization mode. The conjugates PAA-Asp (1) and PAA-Glu (2) could be identified from methanol extracts of leaves of *PtAAS1*-expressing *N. benthamiana*. Base peak chromatograms and extracted ion chormatograms are shown for the characteristic mother ion as well as one characteristic fragment (grey). Mass spectra of the in source fragmentation patterns for previously identified compounds PAA-Asp (1) and PAA-Glu (2) are shown. A representative sample of the total pool of replicates (n=6) was selected for visualization.

**Supplemental Figure 6:**
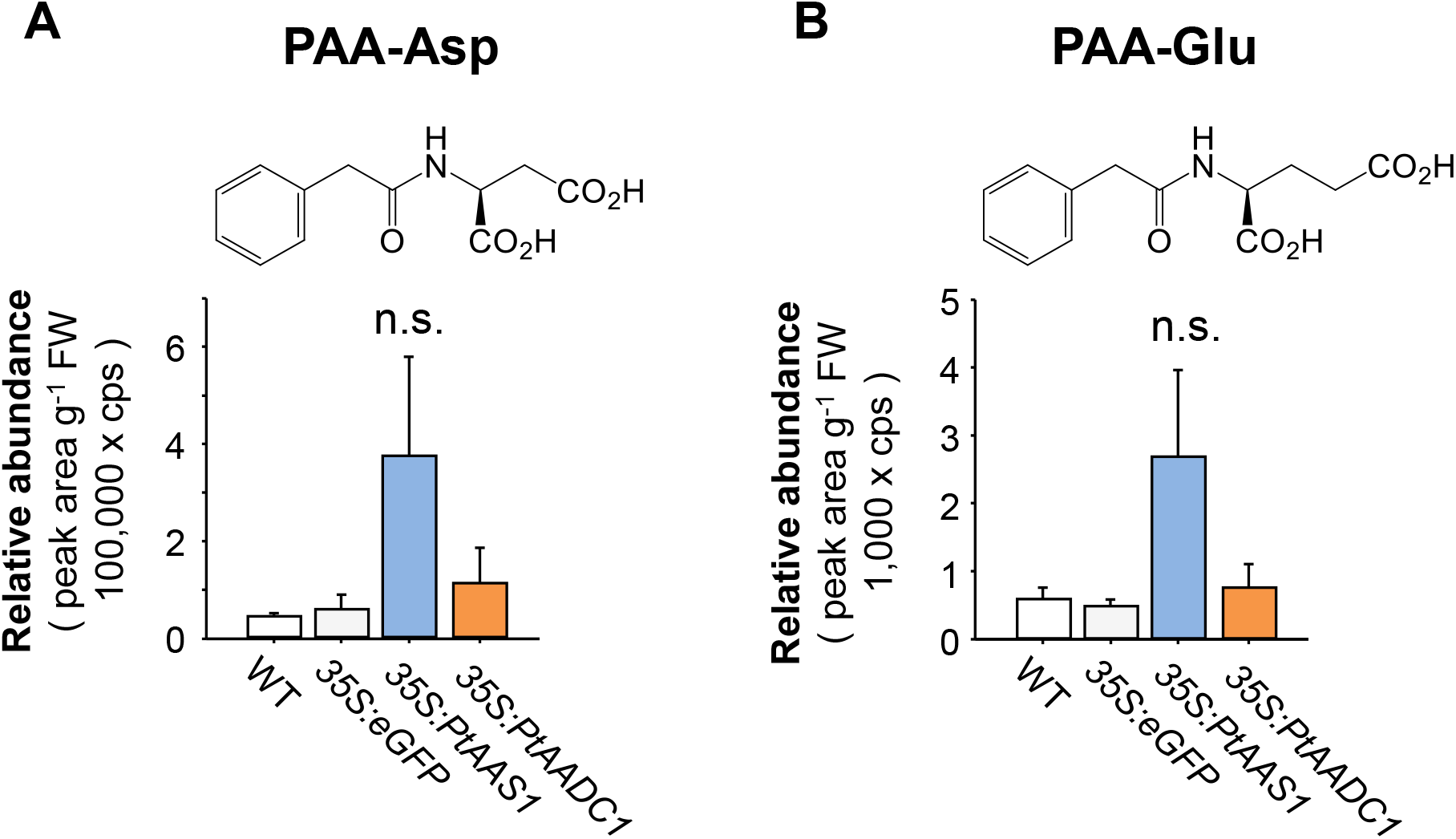
Expression of *PtAAS1* results in unaltered abundance of auxin conjugates PAA-Asp (A) and PAA-Glu (B) in *N. benthamiana* roots. The identified conjugates were analyzed for a characteristic fragmentation via LC-MS/MS. Means + SE are shown (n = 6). FW, fresh weight. n.s., not significant

**Supplemental Figure 7:**
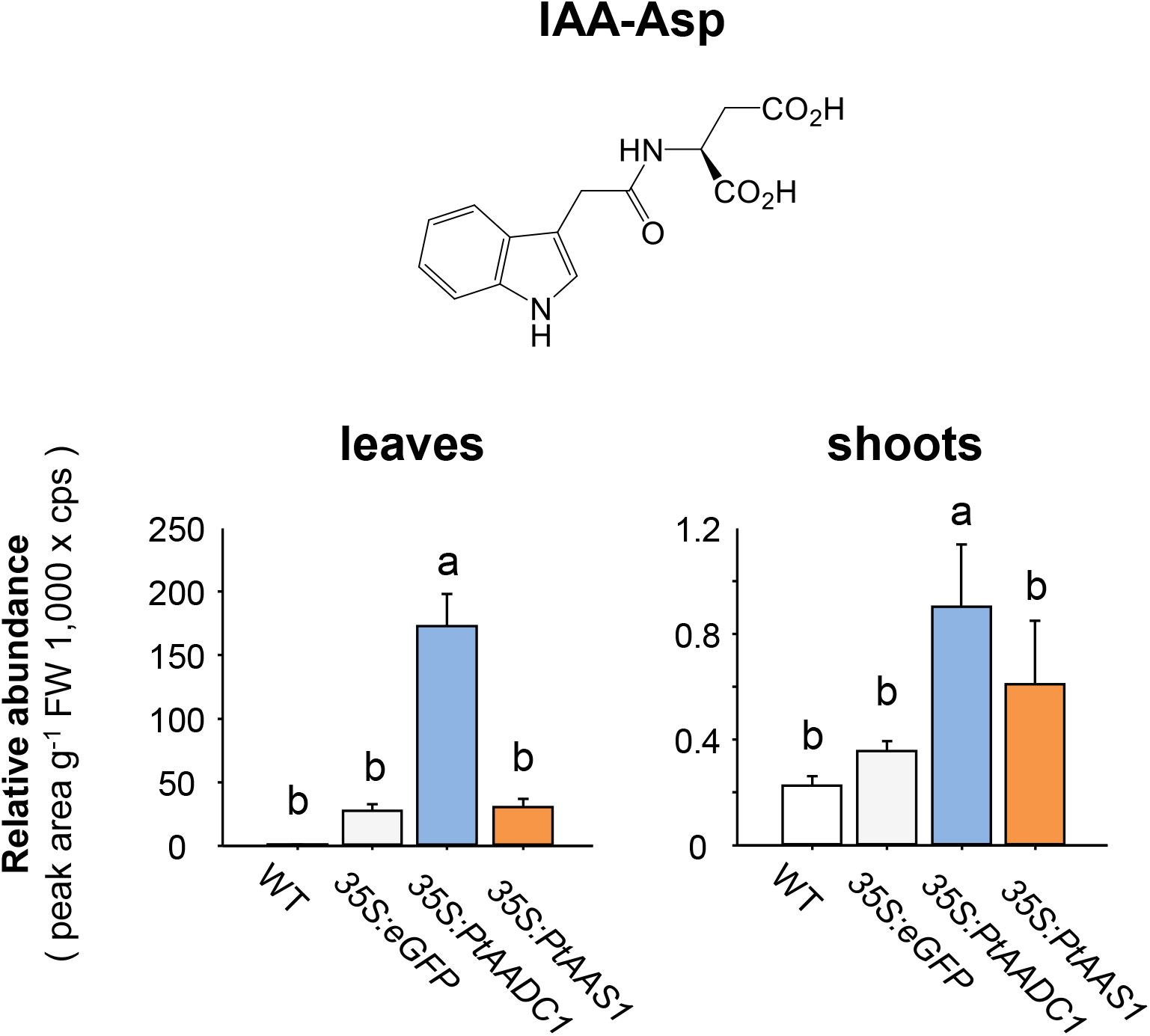
Expression of *PtAAS1* results in increased levels of the auxin conjugate IAA-Asp in *N. benthamiana* shoots and leaves. The identified conjugates were analyzed for a characteristic fragmentation via LC-MS/MS. Relative quantification of the identified conjugates IAA-Asp. Different letters above each box indicate statistically significant differences in Kruskal-Wallis One Way ANOVA and are based on the following Tukey test. IAA-Asp_Shoots_ (H = 15.173, P = 0.002); IAA-Asp_Leaves_ (H = 19.547, P ≤ 0.001). Means + SE are shown (n = 6) FW, fresh weight.

**Supplemental Figure 8:**
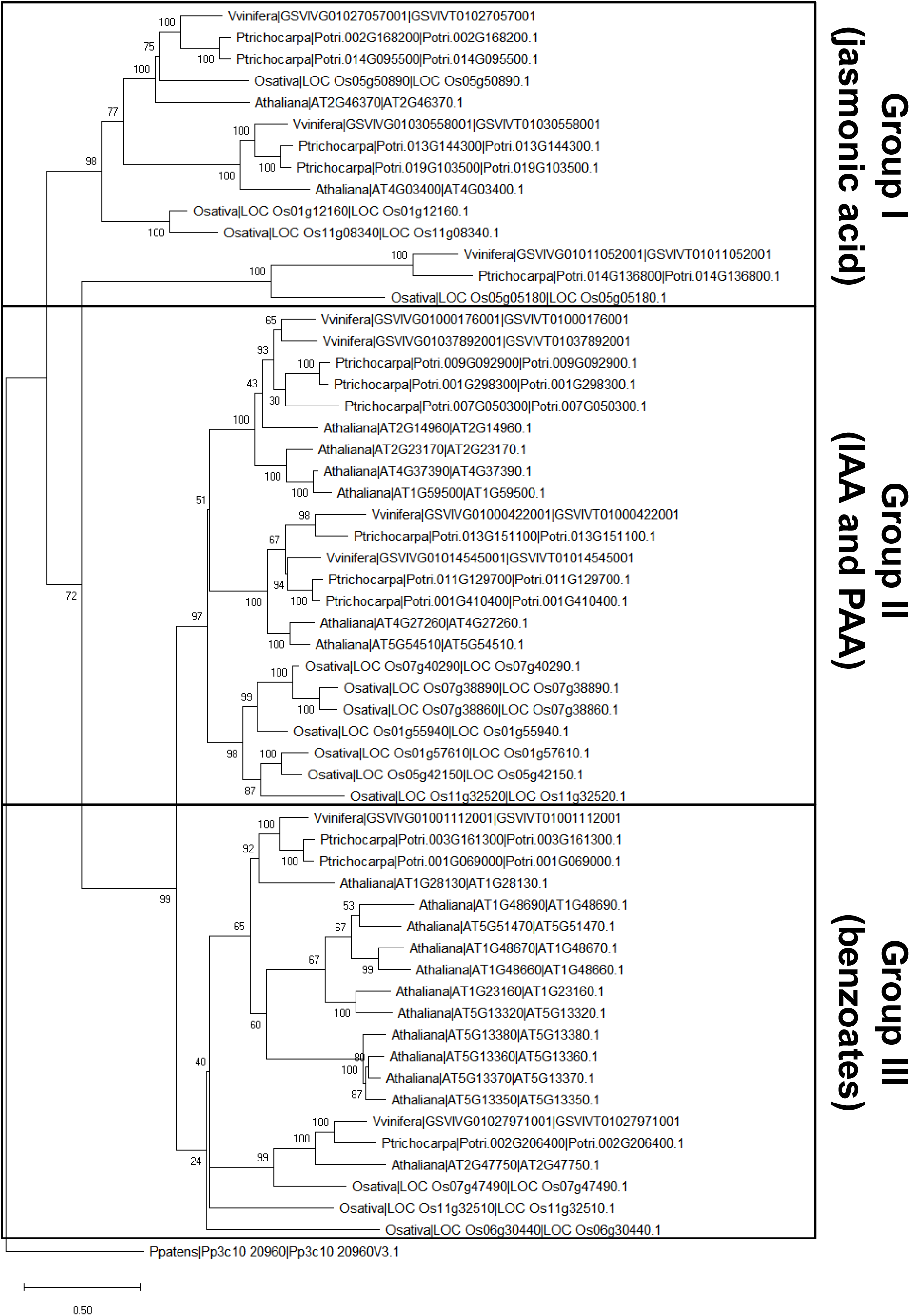
Phylogenetic reconstruction of identified and characterized GH3 auxin-amido synthetases coding sequences. Putative GH3 auxin-amido synthetase sequences from *Populus trichocarpa* and recently identified and characterized GH3 auxin-amido synthetases from *Oryza sativa*, *Arabidopsis thaliana*, and *Vitis vinifera*. Each group I - III is highlighted in rectangles and labeled with characteristic substrates. A putative GH3 from *Physcomitrella patens* served as outgroup. The tree was inferred by using the maximum likelihood method and n = 1,000 replicates for bootstrapping. Bootstrap values are shown next to each node. Relative branch lengths measure the number of substitutions per site.

**Supplemental Figure 9:**
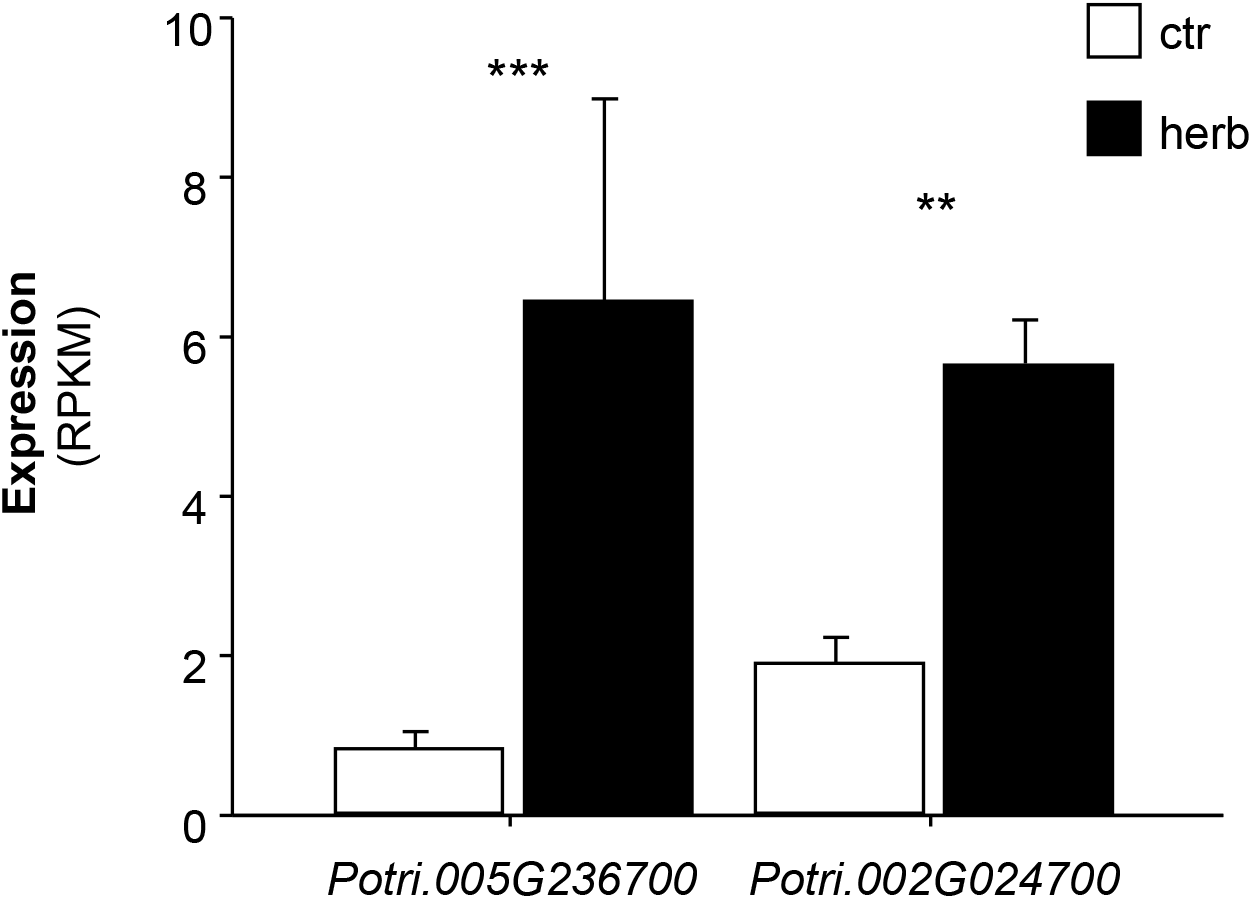
Transcript accumulation of *Aux/IAA* genes in *L. dispar*-damaged and undamaged *Populus trichocarpa* leaves. Gene expression in herbivore-damaged (herb) and undamaged (ctr) leaves was analyzed by Illumina sequencing and mapping the reads to the transcripts of the *P. trichocarpa* genome version v3.0. Expression was normalized to RPKM. Significant differences in EDGE tests are visualized by asterisks. Means + SE are shown (n = 4). *Potri.005G236700* (P = 4.06063E-05, weighted difference (WD) = 8.81922E-06); *Potri.002G024700* (P = 0.005250486, WD = 6.0016E-06).

