## Supplemental infromation for "Heterologous expression of *PtAAS1* reveals the metabolic potential of the common plant metabolite phenylacetaldehyde for auxin synthesis *in planta*"

**Supplemental Table 3**

**HPLC conditions (gradients) used for separation and analysis of plant metabolites.** See also Supplemental Table 2 for MS/MS parameters.

| | **Time** (min) | **Flow** (µl/min) | **Water 0.5% FA** (%) | **Acetonitrile** (%) |
|---|---|---|---|---|
| **Gradient A (auxins sched MRM)** | 0 | 1100 | 95 | 5 |
| | 0.5 | 1100 | 95 | 5 |
| | 6.0 | 1100 | 62.6 | 37.4 |
| | 6.02 | 1100 | 0 | 100 |
| | 7.5 | 1100 | 0 | 100 |
| | 7.6 | 1100 | 95 | 5 |
| **Gradient B (auxin-conjugates)** | 0 | 1100 | 95 | 5 |
| | 0.5 | 1100 | 95 | 5 |
| | 6 | 1100 | 62.6 | 37.4 |
| | 6.02 | 1100 | 20 | 80 |
| | 7.5 | 1100 | 0 | 100 |
| | 9.5 | 1100 | 0 | 100 |
| | 9.52 | 1100 | 95 | 5 |
| **Gradient C (amines and amino acids)** | 0 | 1100 | 97 | 3 |
| | 1 | 1100 | 97 | 3 |
| | 2.7 | 1100 | 0 | 100 |
| | 3 | 1100 | 0 | 100 |
| | 3.1 | 1100 | 97 | 3 |
| | 6 | 1100 | 97 | 3 |

**Supplemental Table 4**

**MS/MS parameters used for LC-MS/MS analysis.** The details of the HPLC gradients indicated in the right column are given in Supplemental Table 2.

| compound | Q1 | Q3 | Scan time (ms) | DP | EP | CE | CXP | mode | HPLC gradient |
|---|---|---|---|---|---|---|---|---|---|
| phenylacetic acid[1] | 134.854 | 91 | 1000 | -30 | -8.5 | -10 | -10 | neg | B |
| PAA-Asp[2] | 250.1 | 132.1 | 20 | -30 | -8.5 | -15 | -10 | neg | A |
| PAA-Glu[2] | 264.173 | 146.1 | 20 | -30 | -8.5 | -15 | -10 | neg | A |
| IAA-Asp[3] | 289 | 132 | 20 | -55 | -9 | -24 | -10 | neg | A |
| 4-OH-PAA[1] | 151.05 | 107.0 | 1000 | -30 | -8.5 | -10 | -10 | neg | B |
| 2-PEG[4] | 329.015 | 45 | 20 | -30 | -5.5 | -24 | 0 | neg | A |
| IAA[1] | 176 | 130 | 10 | 40 | 10 | 19 | 16 | pos | C |
| Phe[4] | 166.2 | 120.2 | 10 | 30 | 6 | 17 | 4 | pos | C |
| Tyr[4] | 182.1 | 136.2 | 10 | 30 | 7 | 17 | 4 | pos | C |
| Trp[4] | 205.2 | 188.1 | 10 | 30 | 4.5 | 13 | 6 | pos | C |
| PEA[5] | 122.01 | 105 | 10 | 30 | 4 | 15 | 4 | pos | C |
| TyrA[5] | 138.01 | 121 | 10 | 30 | 4 | 15 | 4 | pos | C |
| TrpA[5] | 181.2 | 144.2 | 10 | 30 | 8 | 17 | 4 | pos | C |

**Supplemental Table 5**

**Compounds used as standards for LC-MS/MS quantification.**

| Compound | CAS | Supplier |
|---|---|---|
| L-phenylalanine | 63-91-2 | Duchefa |
| L-tyrosine | 60-18-4 | Duchefa |
| L-tryptophan | 73-22-3 | SigmaAldrich |
| 2-phenylethylamine | 156-28-5 | Santa Cruz |
| tyramine | 60-19-5 | Roth |
| tryptamine | 61-54-1 | AppliChem |
| 2-phenylethyl-β-D-glucopyranoside | 18997-54-1 | Toronto Research Chemicals |
| Indole-3-acetic acid | 87-51-4 | Duchefa |
| 4-hydroxy phenylacetic acid | 156-38-7 | Acros |
| phenylacetic acid | 103-82-2 | Aldrich |
